# Somatosensory innervation of healthy human oral tissues

**DOI:** 10.1101/2021.02.03.429664

**Authors:** Yalda Moayedi, Stephanie Michlig, Mark Park, Alia Koch, Ellen A Lumpkin

## Abstract

The oral somatosensory system relays essential information about mechanical stimuli to enable oral functions such as feeding and speech. The neurochemical and anatomical diversity of sensory neurons across oral cavity sites have not been systematically compared. To address this gap, we analyzed healthy human tongue and hard palate innervation. Biopsies were collected from 12 volunteers and underwent multiplex fluorescent immunohistochemistry (≥2 specimens per marker/structure). Afferents were analyzed for markers of neurons (βIII tubulin), myelinated afferents (neurofilament heavy, NFH), and Merkel cells and taste cells (keratin 20, K20). Hard-palate innervation included Meissner’s corpuscles, glomerular endings, Merkel cell-neurite complexes, and free nerve endings. The organization of these somatosensory endings is reminiscent of fingertips, suggesting that the hard palate is equipped with a rich repertoire of sensory neurons for pressure sensing and spatial localization of mechanical inputs, which are essential for speech production and feeding. Likewise, the tongue is innervated by afferents that impart it with exquisite acuity and detection of moving stimuli that support flavor construction and speech. Filiform papillae contain end bulbs of Krause, as well as endings that have not been previously reported, including subepithelial neuronal densities, and NFH+ neurons innervating basal epithelium. Fungiform papillae had Meissner’s corpuscles and densities of NFH+ intraepithelial neurons surrounding taste buds. The differing compositions of sensory endings within filiform and fungiform papillae suggest that these structures have distinct roles in mechanosensation. Collectively, this study has identified previously undescribed afferent endings in human oral tissues and provides an anatomical framework for understanding oral mechanosensory functions.

## Introduction

Mechanosensation in the oral cavity has long been hypothesized to provide inputs for essential functions including speech production, flavor construction, and coordination of feeding mechanics. The importance of oral somatosensation is evident by the large area of primary somatosensory cortex devoted to the oral cavity in humans (Tamura et al., 2008). In production of speech, mechanosensory feedback from the tip of the tongue and hard palate is essential for articulation of vowels and sibilants (Hickok, 2012; Niemi et al., 2006; Niemi et al., 2002). The sensation of food textures can drive nutrient choice, perceived fullness after consumption, and overall caloric intake, suggesting that mechanosensation plays an important role in feeding (McCrickerd & Forde, 2016). During ingestion, the size and consistency of a food bolus sensed through mechanosensory afferents are believed to guide phases of ingestion, mastication, and swallowing (Inoue et al., 1989; Moore et al., 2014; Thexton et al., 1980; Travers & Norgren, 1986). Despite the importance of mechanosensation in nutrient intake and social interactions, the cellular substrates of touch in the oral cavity are unclear.

In skin, mechanical stimulus features are transduced by an array of mechanosensory cells and neurons (Zimmerman et al., 2014). These mechanosensory neurons have distinct peripheral morphologies and response properties, which allow them to encode different perceptual features of touch (Moayedi et al., 2015). For example, Meissner’s corpuscles are rapidly adapting (RA) afferents that respond best to low frequency vibrations and encode a sensation of flutter (Mountcastle et al., 1967). Merkel cell-neurite complexes, on the other hand, fire throughout a sustained stimulus generating the slowly adapting type I (SAI) response, and encode textures and edges (Iggo & Muir, 1969; Johnson et al., 2000). The distinct peripheral morphologies of mechanosensory neurons along with their accessory cells and structures, collectively called end organs, can be identified via histochemical methods (Cobo et al., 2021). The composition of these end organs varies over the surface of the body, and the cadre of sensory endings determine the range of sensations experienced by a particular anatomical location.

Compared with skin innervation, less is known about innervation of oral cavity structures. A previous report identified the complement of anatomically and neurochemically distinct neuronal endings in mouse oral cavity using transgenic reporters, neurochemical markers and quantitative histomorphometry (Moayedi et al., 2018). The mouse tongue is equipped with an array of NFH-positive putative mechanoreceptors that express the mechanosensory channel, Piezo2. These included end bulbs of Krause innervating individual filiform papillae and neuronal afferents innervating the epithelium surrounding taste buds. The mouse hard palate is densely populated with three classes of sensory afferents organized in discrete patterns that resemble that of glabrous skin. These include Merkel cell-neurite complexes, Meissner’s corpuscles, and glomerular endings-a type of non-lamellar sensory corpuscle composed of a mixture of entangled neuronal endings and Schwann cells (Byers & Yeh, 1984). The diverse array and high density of mechanosensory afferents embedded in mouse oral surfaces suggest that the tongue and hard palate are tuned to detect distinct features of mechanical stimulation with high tactile acuity.

Human oral surfaces display rich mechanosensitivity. In particular, the tip of the tongue has comparable acuity and sensitivity to that of the fingertip (Miles et al., 2018; Van Boven & Johnson, 1994). Despite the importance of sensory inputs for feeding, few studies have investigated the distribution of somatosensory endings in human healthy oral epithelia (Hilliges et al., 1996; Marlow et al., 1965; Ramieri et al., 1992). Here, we present a systematic analysis of the complement of somatosensory neurons by using a panel of neurochemical markers to investigate innervation of biopsies from defined regions of tongue and hard palate.

## Methods

### Tissues from donation

Biopsies of the hard palate and tongue were obtained from three male anatomical donors. Donations occurred through the Anatomical Donor Program of Columbia University’s Vagelos College of Physicians and Surgeons’. Punch biopsies (4-mm diameter) were collected from two sites: the front of tongue or palate rugae.

### Human healthy volunteer tissue collection

All human studies were approved by the Institutional Review Board of Columbia University. Adult volunteers (18-50 years old) were eligible for enrollment in this study (**Table 1**). Exclusion criteria included: current infection, pain, injury or other cutaneous abnormality in the oral cavity that the investigator feels may interfere with study interpretation or safely performing a biopsy; current use of anticoagulants (e.g. aspirin, coumadin, NSAIDs), history of a bleeding disorder, history of keloidal or hypertrophic scarring, oral cancer, neurological diseases that may affect innervation of the oral cavity, known or suspected abnormalities of epithelial innervation in the area to be biopsied, any known or suspected medical or psychological conditions that may affect the subjects ability to properly consent to participation or to follow instructions for wound care, active medical conditions that the investigator deems may affect the patient’s risk of infection or ability to heal normally after the biopsy.

**Table 1.** Collected biopsies.

| Site of Biopsy | Age | Sex |
| --- | --- | --- |
| Palate Back | 27 | M |
| Palate Back | 31 | F |
| Palate Rugae | 40 | F |
| Palate Rugae | 30 | F |
| Palate Rugae | 43 | F |
| Tongue Back | 32 | F |
| Tongue Back | 45 | F |
| Tongue Front | 50 | F |
| Tongue Front | 38 | F |
| Tongue Front | 43 | F |
| Tongue Front | 33 | F |
| Tongue Front | 27 | F |

### Informed consent

Written informed consent was obtained for this study by personnel from all patients prior to any protocol-specific procedures. The study was conducted in accordance with the Food and Drug Administration (FDA) approved revision of the Declaration of Helsinki, current FDA regulations, and International Conference on Harmonization (ICH) guidelines.

### Tissue collection

One-site was biopsied from each participant, either the front of tongue, middle of tongue, palate rugae, or post-rugal field of the hard palate (**Figure 1**). Biopsy site was anesthetized with an injection of 2% lidocaine with epinephrine (1:100,000). After approximately five minutes, the clinician used a 4-mm punch to collect the biopsy. The punch was oriented to be perpendicular to the surface epithelium. The punch went down to the submucosal layer. Non-traumatic forceps were used to gently grasp the specimen, pulling upward to remove the core and reveal the submucosal layer. Scissors were used to cut the tissue free from the underlying submucosal layer, if needed. The specimen was placed immediately into phosphate buffered saline (PBS). Once the specimen was removed, pressure was applied to the biopsy site with sterile 2X2 gauze. The biopsy site was then closed with several simple resorbable interrupted sutures, if indicated. Additional gauze was applied to the site intraorally and subjects were instructed in wound care and advised to call the research unit if they had any concerning signs or symptoms during healing. Compensation was given after biopsies were taken. Age and sex information about specimens collected in this study are described in **Table 1.**

**Figure 1.**
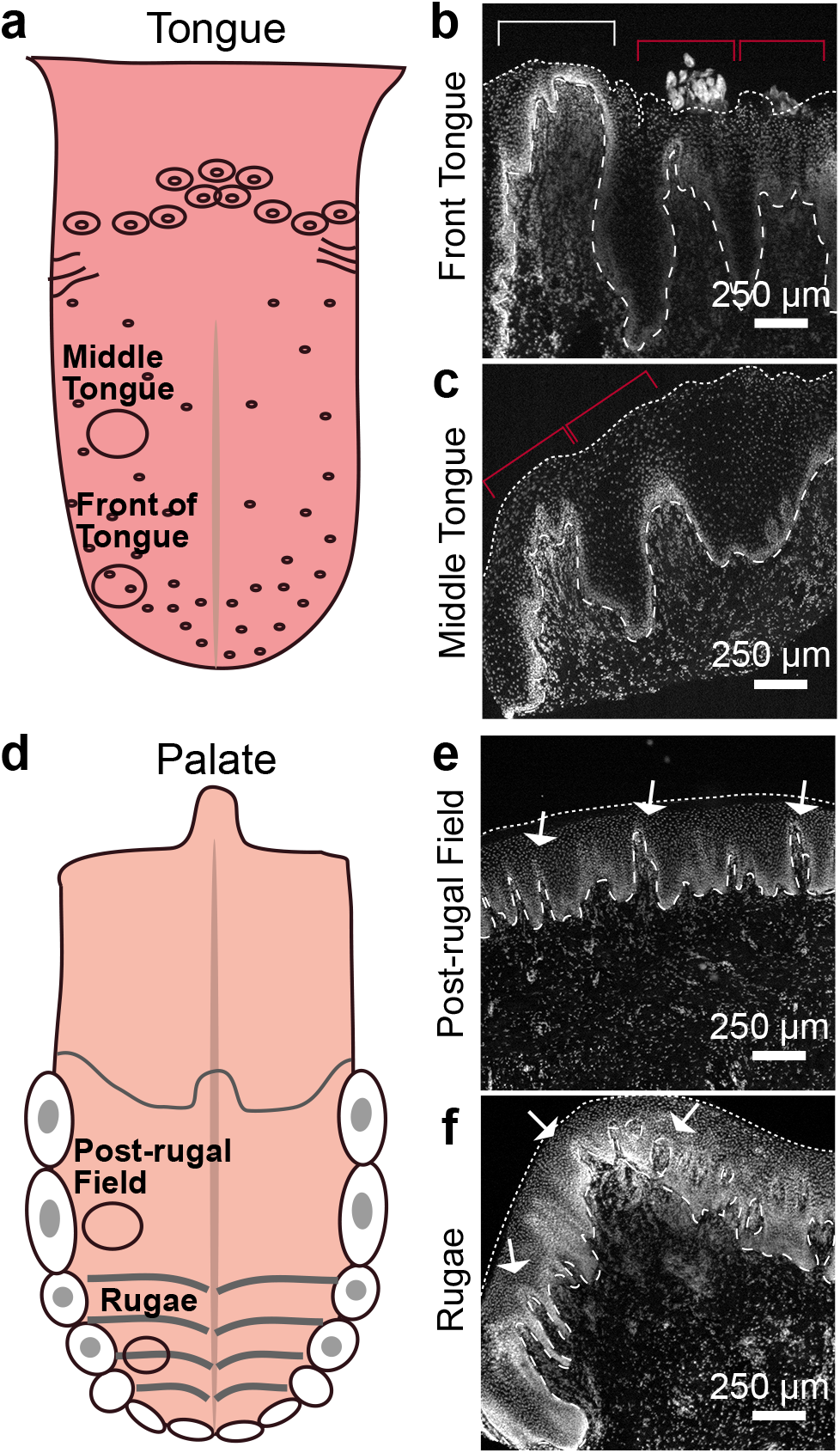
Collection sites and regional ultrastructure. Schematics showing collection sites on tongue and hard palate are shown (left) and sample ultrastructural images of DAPI staining are shown for each site (right). Dotted lines indicate epithelial surface. Dashed lines indicate epithelial-lamina propria border. **a.** Biopsies were collected from the middle and front of tongues. **b.** Front of tongue biopsies contained fungiform papillae (white bracket) and filiform papillae (red brackets). **c.** Middle tongue biopsies primarily consisted of filiform papillae (red brackets). **d.** Hard palate biopsies were collected from the post-rugal field and palate rugae. **e.** Post-rugal field has a flat surface with lamina propria pits jutting into epithelial layer (white arrows). **f.** Biopsies from rugae have a curved surface with lamina propria pits similar to the post-rugal field (white arrows).

### Tissue analysis

After collection, tissues were photographed, embedded in TissueTec OCT, and flash frozen. Tissue samples were sectioned at a thickness of 25 μm on gelatinized slides. Slides were incubated for 30 min at 37°C then fixed with 4% paraformaldehyde for 10 min followed by five washes in PBS. Slides were blocked in PBS with 0.03% triton-X 100 (PBST) and 5% normal goat serum (hybridization buffer). Sections were then hybridized overnight with primary antibodies mixed in hybridization buffer at 4°C. Primary antibodies used in this study are described in **Table 2**. The following day, slides were washed thrice in PBS and then incubated with secondary antibody in hybridization buffer for 1–2 h. Secondary antibodies used in this study are described in **Table 3**. Slides were then washed five times in PBS and mounted in Fluoromount-G with DAPI. Specimens were imaged with a laser scanning confocal microscope equipped with 40X (NA 1.3) and 20X (NA 0.8) lenses. Images were processed using ImageJ and Adobe Photoshop. Adjustments for brightness, contrast, and threshold were made in Adobe Photoshop and applied to the entire image.

**Table 2.** Primary Antibodies.

| Antibody | Cells marked | Source | Dilution | Catalog # | Lot # | RRID |
| --- | --- | --- | --- | --- | --- | --- |
| Rabbit anti-βIII tubulin | All neurons | Abcam | 1:3000 | ab18207 | GR3221401-3 | AB_444319 |
| Mouse anti-Keratin 20 | Taste cells, Merkel cells | Abcam | 1:100 | ab854 | GR157163-5 | AB_2133708 |
| Chicken anti-Neurofilament-Heavy | Myelinated Neurons | Abcam | 1:5000 | ab4680 | GR310109-11 | AB_304560 |

**Table 3:** Secondary Antibodies.

| Antibody | Source | Dilution | Catalog # | RRID |
| --- | --- | --- | --- | --- |
| Alexa 488 conjugated goat anti-rabbit | Thermo-Fisher | 1:1000 | A-11008 | AB_143165 |
| Alexa 594 conjugated goat anti-rabbit | Thermo-Fisher | 1:1000 | A-11012 | AB_2534079 |
| Alexa 594 conjugated goat anti-mouse | Thermo-Fisher | 1:1000 | A-11032 | AB_2534091 |
| Alexa 488 conjugated goat anti-mouse | Thermo-Fisher | 1:1000 | A11029 | AB_138404 |
| Alexa 647 conjugated goat anti-chicken | Thermo-Fisher | 1:1000 | A-21449 | AB_2535866 |

## Results

To analyze the diversity of somatosensory neurons innervating human oral cavity, we investigated anatomical and neurochemical features of sensory afferents in human palate and tongue specimens. We performed immunolabelling of tissue sections with markers for all neurons (βIII-tubulin, βIII), myelinated neuronal afferents (Neurofilament Heavy, NFH), and Merkel cells/taste cells (Keratin 20, K20). This biomarker panel allows for distinction of unmyelinated (βIII+, NFH-) and myelinated (βIII+, NFH+) neuronal afferents, representing putative nociceptors and mechanoreceptors, respectively. Tissue collection locations were based on our previous findings in mice in order to provide a systemic analysis of neuronal diversity in these regions and to allow for cross-species comparisons. Specimens were collected from four sites in the oral cavity: middle tongue, front of tongue, hard palate post-rugal field, hard palate rugae (**Figure 1**). Sections stained with the nuclear marker DAPI revealed the structure of the tissue. Middle tongue (**Figure 1a-b, red arrows**) contained abundant filiform papillae, the non-taste papillae that cover most of the tongue. Specimens from the front of the tongue contained filiform papillae and fungiform papillae, the taste papillae that speckle the anterior two-thirds of the tongue. In pilot experiments with the taste-cell marker K20, we observed that fungiform papillae have greater height than filiform papilla, a thinner epithelial layer, and the presence of a larger lamina propria core; therefore, these three anatomical hallmarks were used to identify fungiform papillae in sections (**Figure 1a, c, white arrow**). Filiform papillae, on the other hand, were relatively short with flatter morphology and thinner lamina propria cores compared with fungiform papillae. The post-rugal field had a flat surface lined with epithelial pegs (**Figure 1d-e**, **white arrows**). Hard-palate rugae had a curved epithelial surface and abundant epithelial pegs. (**Figure 1d, f, white arrows**). These macroscopic features were used throughout the study to orient neurochemical identification of innervation.

We began neurochemical analysis by performing pilot studies in tongue specimens from anatomical tissue donations (**Figure 2**). Human fungiform papillae were innervated by a dense meshwork of NFH+ and NFH-afferents both below and within the taste bud (**Figure 2a**). These neurons include taste afferents with cell bodies in the geniculate ganglion. NFH+ and NFH-neuronal afferents were also found to innervate the epithelia surrounding the taste buds (**Figure 2a, white and red arrows**). These afferents would likely include neurons transducing chemical and thermal stimuli as well as mechanosensory neurons. In addition, putative corpuscular endings with morphology similar to a Meissner’s corpuscle was identified in two specimens, residing deep within fungiform papillae (**Figure 2b)**. Meissner’s corpuscles had typical oblong morphology consisting of a myelinated (NFH+) neuron spiraling apically, with the long axis of the corpuscle perpendicular to the epithelial surface. In filiform papillae, end bulbs of Krause were identified as the predominant type of end organ (**Figure 2c, white arrows**). These ending types were identified via immunohistochemistry as myelinated afferents that take a convoluted path within a spherical corpuscle, compared with Meissner’s corpuscles where endings take helical turns within an oblong corpuscles (Chouchkov, 1973). Multiple end bulbs of Krause populated a single filiform papilla **(Figure 2c, white arrows**), ranging from 1–5 total end bulbs within a single 25-μm section of a filiform papillae (n=4 papillae analyzed). Short extensions of NFH+ afferents into epithelium were occasionally identified in specimens (**Figure 2c, blue arrow**). In addition to myelinated afferents, occasional unmyelinated (NFH-, βIII+) afferents were identified coursing through the lamina propria towards the apical portions of filiform papillae (**Figure 2c, red arrow**). We also identified sparse K20+ cells with Merkel cell-like morphology at the epithelial-lamina propria (**Figure 2d, white arrow**). These K20+ cell clusters were identified in three papillae, and contained between one and three total cells. Notably, neuronal afferents were not observed near these K20+ cells, indicating that they might not be innervated. Thus, the human tongue samples displayed rich innervation with a variety of end-organ types present.

**Figure 2.**
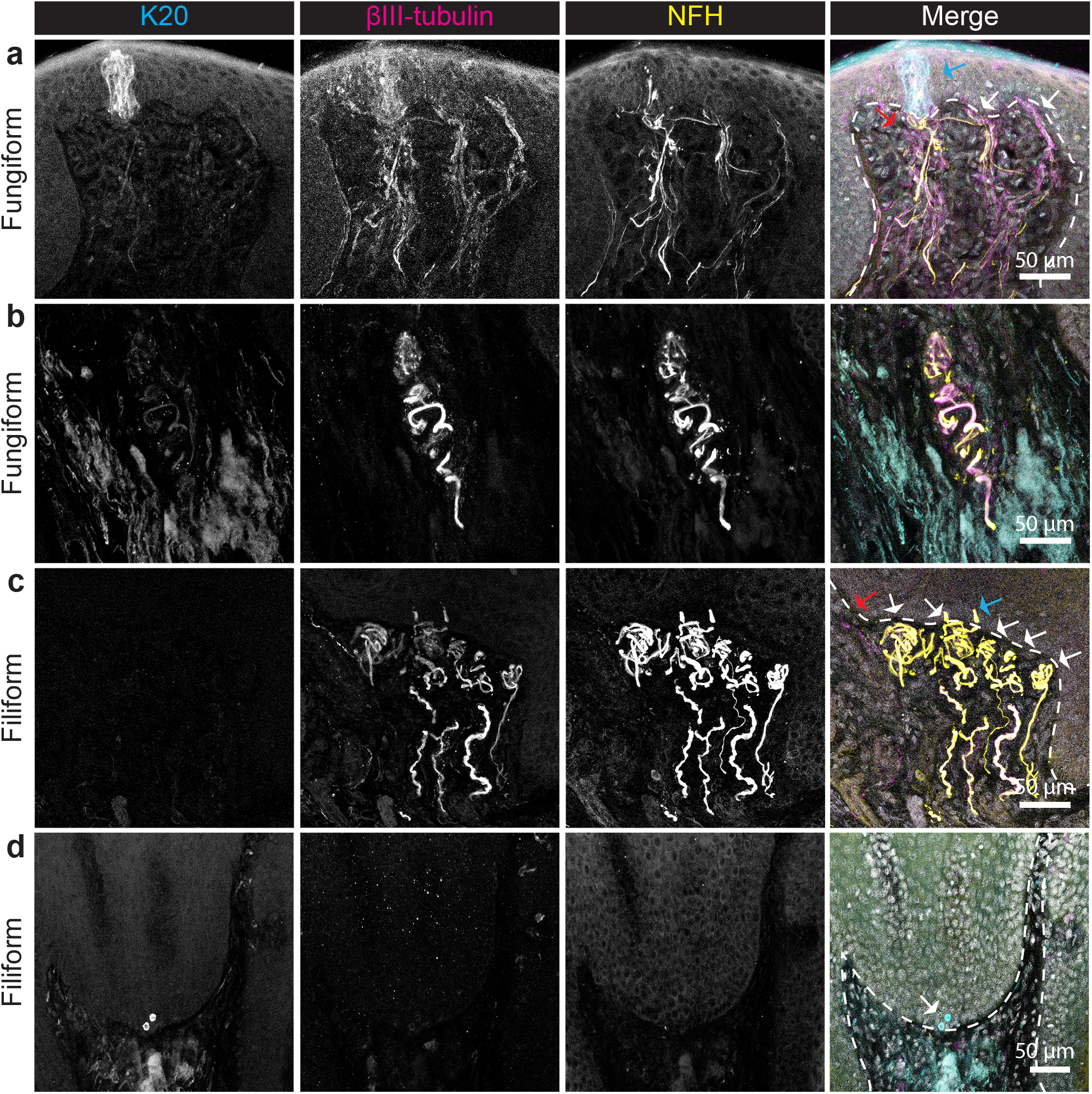
Putative mechanosensory endings are visualized via an immunohistochemical panel in tongue biopsies. Tongue biopsies from donated specimens were sectioned and stained with markers for taste cells and Merkel cells (K20, left column), all neurons (βIII-tubulin, second column), and myelinated neurons (NFH, third column). Epithelial-lamina propria boundary is demarcated by dashed lines. **a.** Fungiform papillae contain taste buds visualized by K20 staining. NFH+ and NFH-fibers innervate subepithelial space below taste buds (blue arrow), adjacent to taste buds (white arrows), and within taste buds (red arrows). **b.** Meissner corpuscles were identified within fungiform papillae, distinguished by intertwined NFH+ and NFH-nerve fibers. **c.** Filiform papillae contained multiple end bulbs of Krause (white arrows), with prominent NFH+ afferents. Blue arrow denotes NFH+ fibers innervating epithelium. Red arrow indicates NFH-fibers coursing through lamina propria. **d.** Occasional clusters of Merkel cells (white arrows) were identified in epithelium of filiform papillae at lamina propria junction.

Innervation of the human hard palate was next investigated in anatomical donor specimens (**Figure 3**). The human hard palate was lined with innervated epithelial pegs. We identified occasional glomerular type endings in hard palate rugae, (**Figure 3a, white arrow**). Glomerular endings were identified by immunostaining through the presence of meandering neuronal afferents forming bundles in the apical portions of hard palate rugae lamina propria. These endings were associated with ultraterminals (**Figure 3a, blue arrow**), identified as afferents extending from the tip of the corpuscle into the rugae epithelium (Gairns, 1954). Meissner-like endings were also found in palates (**Figure 3b, white arrow**). These were similar in structure to those found in tongue, and resided in apical regions of lamina propria of epithelial pegs. Sparse Merkel cells were identified in the base of epithelium (**Figure 3b, blue arrow**). Overall, we found that specimens acquired through anatomical donation provided a useful overview of innervation; however, the quality of immunohistochemical staining was irregular with some areas yielding no immunofluorescence, and some afferents displaying patchy immunofluorescence. This could be due to antigen degradation before fixation, health of the donor, uncontrolled timing between donation and collection of specimens, and inconsistent fixation of epithelia. Collectively, inconsistencies in immunofluorescence indicated that histology of donated tissue specimens might yield an incomplete representation of innervation of oral tissues. Likewise, collection of adjacent sites to pathological tissue, which is another common source of human specimens for histology, could have abnormalities in structure due to the underlying pathology. To overcome these limitations, we next collected specimens from volunteers for analysis of oral innervation in healthy adults.

**Figure 3.**
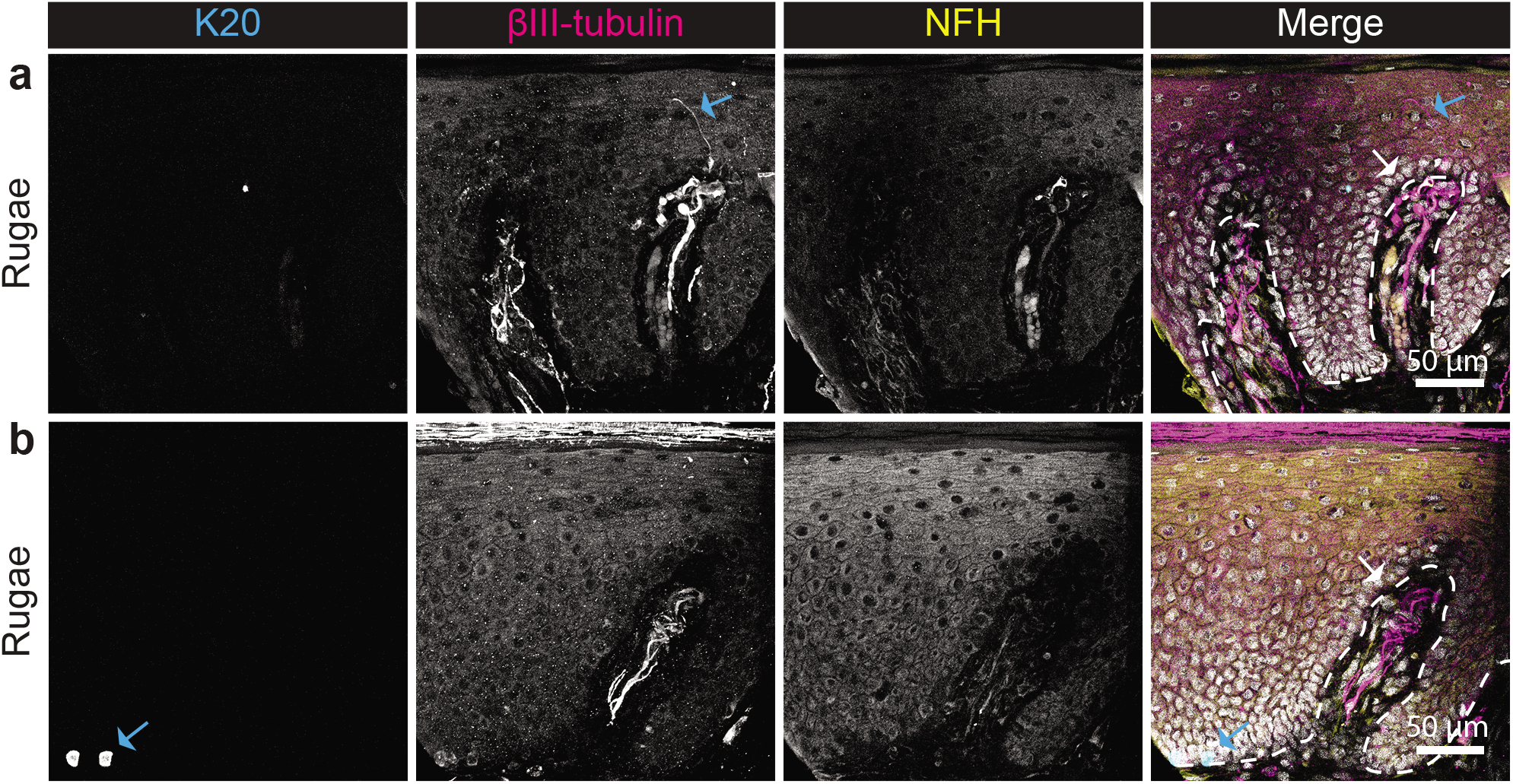
Somatosensory endings can be distinguished in donated hard palate specimens through immunohistochemistry. Palate rugae specimens from donated tissue were sectioned and stained with markers for taste cells and Merkel cells (K20, left column), all neurons (βIII-tubulin, second column), and myelinated neurons (NFH, third column). Epithelial-lamina propria boundary is demarcated by dashed lines. **a.** Glomerular endings (white arrow) with ultraterminals (blue arrow) were identified. **b.** Meissner corpuscles were identified in hard palate biopsies (white arrow), as well as clusters of Merkel cells (blue arrow).

Fungiform papilla from biopsies collected from the front of the tongue were analyzed first (**Figure 4**). Similar to donated specimens, taste buds were identified with K20+ staining, and resided in the apical epithelium of fungiform papillae. One to two taste buds were identified in 25-μm sections of fungiform papillae. Taste buds were associated with dense NFH+ and NFH-afferent bundles in the underlying lamina propria (**Figure 4a, blue arrow**). These fibers include taste afferents innervating taste cells. NFH+ afferents were identified within the taste bud, similar to what has been described in mice (Moayedi et al., 2018). Neuronal afferents innervated the epithelium surrounding taste buds (**Figure 4a, white arrows**). Higher magnification images show that both NFH+ and NFH-neuronal afferents innervate the epithelium adjacent to taste buds (**Figure 4b, white arrow and red arrow, respectively).** NFH-afferents likely include nociceptors responsive to thermal and chemical stimulation, whereas NFH+ neurons are likely to be predominantly mechanoreceptors. A Meissner’s corpuscle was also identified within a fungiform papilla in healthy human tissue (**Figure 4c, white arrow**).

**Figure 4.**
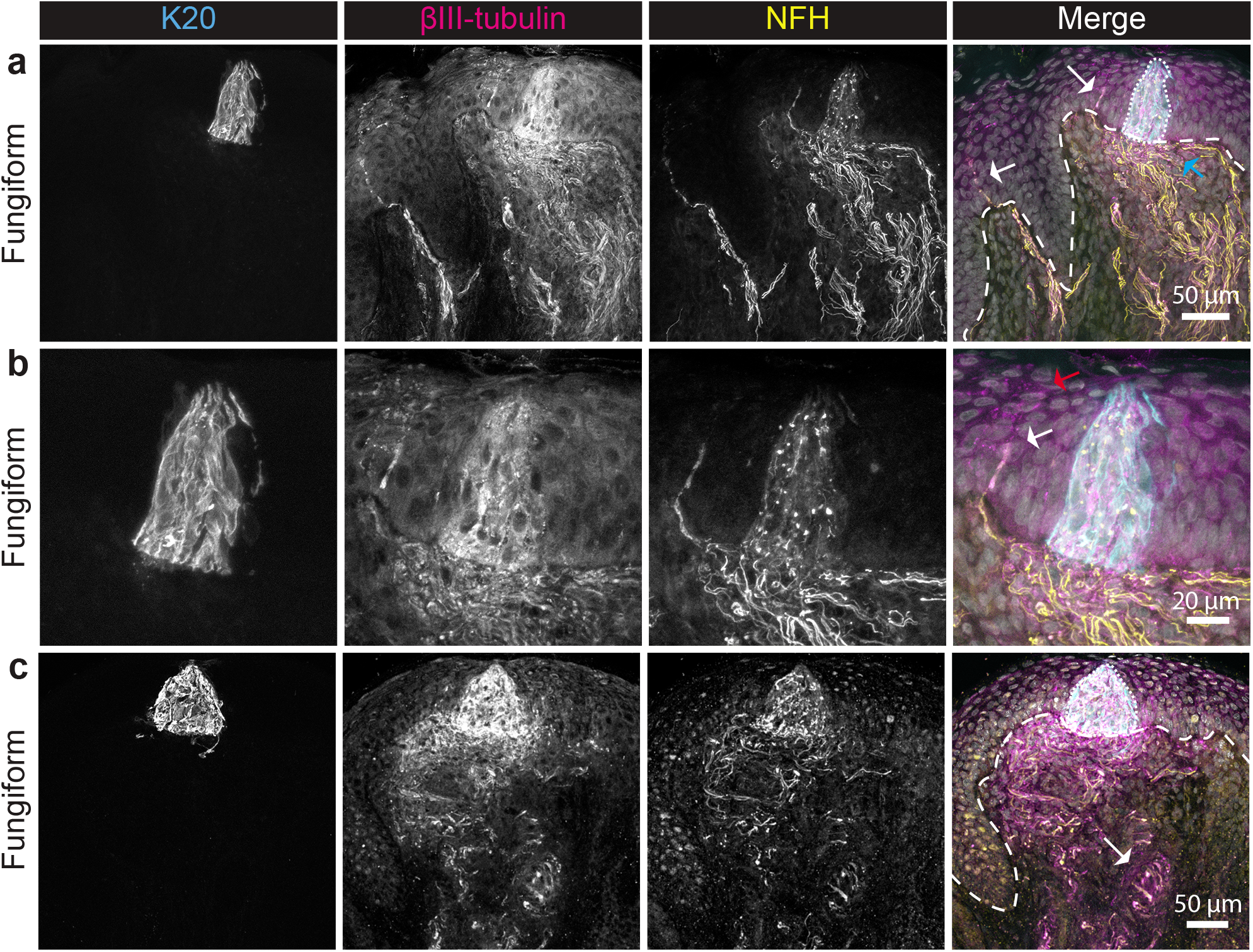
Fungiform papillae are innervated by NFH+ endings terminating in epithelia and Meissner corpuscles within the lamina propria. Tongue biopsies from healthy donors were sectioned and stained with markers for taste cells and Merkel cells (K20, left column), all neurons (βIII-tubulin, second column), and myelinated neurons (NFH, third column). Epithelial-lamina propria boundary is demarcated by dashed lines. **a.** Taste buds were visualized in fungiform papillae through K20 immunoreactivity (dotted line). Dense innervation of NFH+ and NFH-neurons was localized below the taste bud (blue arrow. Neuronal afferents were also identified in the epithelium (white arrows). **b.** A higher magnification view of the taste bud revealed both NFH+ (white arrow) and NFH-(red arrow) afferents within and around the taste bud. **c.** Similar to A, neuronal afferents densely innervated the lamina propria, epithelium, and taste bud of the fungiform papilla. A putative Meissner’s corpuscle was found within this fungiform papilla (white arrow).

In filiform papillae, similar afferent types were observed in donated and living donor specimens. We found end bulbs of Krause within filiform papillae (**Figure 5a, white arrows**), as well as unmyelinated fibers coursing into apical lamina propria (**Figure 5a, red arrow**). A higher magnification view of an end bulb of Krause (**Figure 5b**) shows both myelinated (**white arrow**) and unmyelinated (**red arrow**) afferents within the bulb. Sparse K20+ cells whose morphology resembled Merkel cells within filiform papillae were also observed (**Figure 5c, blue arrow**). As in donated specimens, these cells were not found to be nearby neuronal afferents, indicating that they might not send sensory information to the nervous system. Occasional myelinated afferents contacting the epithelial-lamina propria border were identified in nearby regions (**Figure 5c, white arrow**). We also identified occasional bundles of NFH+ endings terminating just below the epithelial-lamina propria border (**Figure 5d, white arrow**, N=2 specimens), which we term subepithelial neuronal densities. The composition of myelinated endings and the organized subepithelial end-organ structure suggest that these endings may be mechanosensory, and resemble similar mucocutaneous endings described by Marlow and colleagues (Marlow et al., 1965).

**Figure 5.**
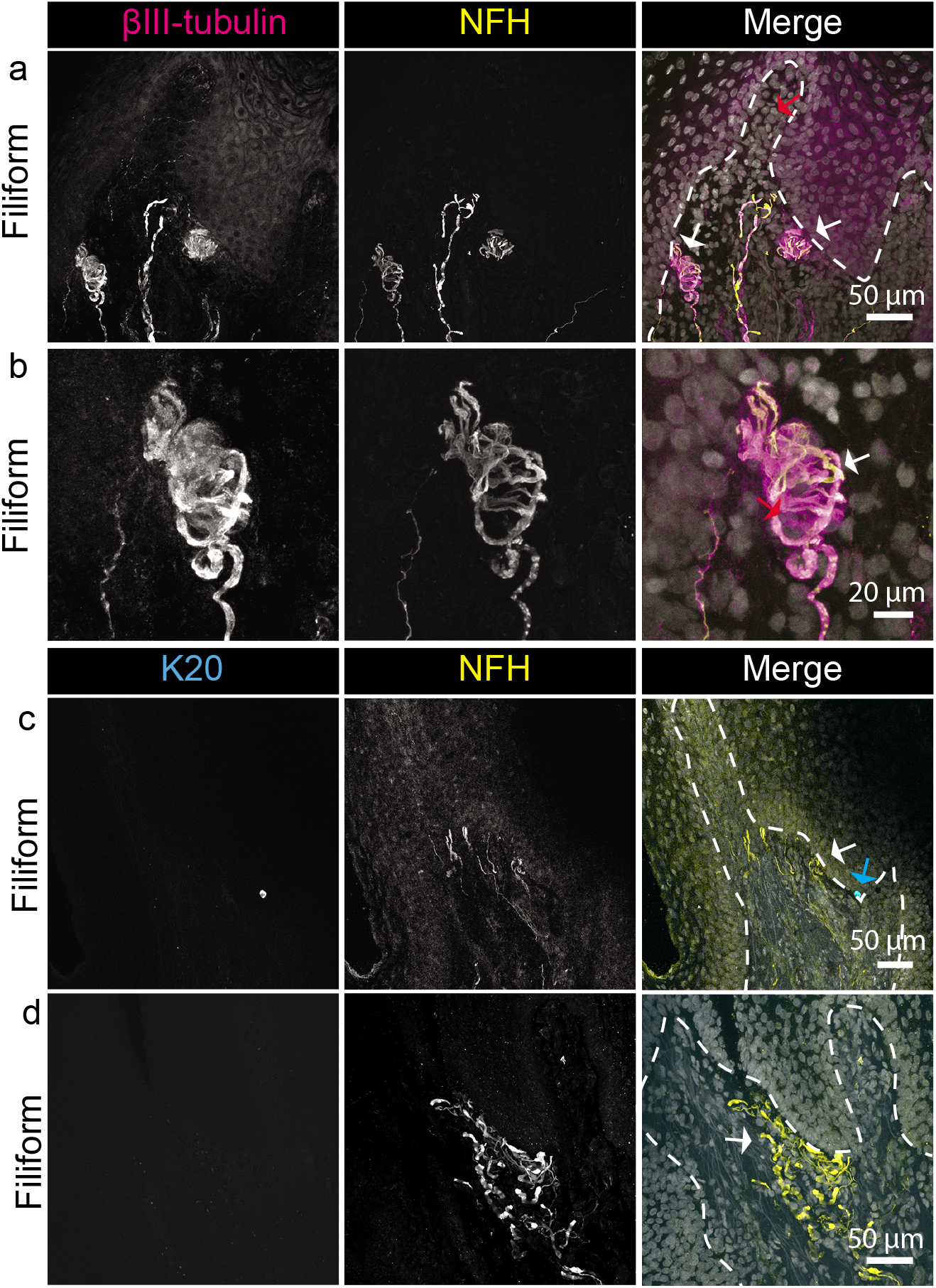
Filiform papillae are innervated by end bulbs of Krause, Merkel cells, subepithelial NFH+ afferents and NFH-neuronal afferents. Tongue biopsies from healthy donors were sectioned and stained with markers for all neurons (βIII-tubulin, left column, top), taste cells and Merkel cells (K20, left column, bottom), and myelinated neurons (NFH, second column). Epithelial-lamina propria boundary is demarcated by dashed lines. **a.** End bulbs of Krause innervated the lamina propria core of filiform papillae (white arrows) visualized by NFH and βIII staining. NFH-βIII+ fibers were also present in the papillary core (red arrow). **b.** High magnification view of End bulbs of Krause in **a.** showing both NFH+ (white arrow) and NFH-(red arrow) fibers. **c.** Filiform papilla subepithelial myelinated afferents (white arrow) and a single Merkel cell (blue arrow). **d.** A filiform papilla showing dense NFH+ neurons coalescing together in the subepithelial space (white arrow).

Hard palates were investigated next (**Figure 6**). Neuronal endings were often found within lamina propria jutting up between epithelial pegs (**Figure 6a, blue arrow**) of hard-palate rugae. Large clusters of Merkel cells were found at the base of epithelial pegs in hard palate rugae (**Figure 6a, white arrow**). In a higher magnification view, Merkel cells were found to be contacted by NFH+ afferents (**Figure 6b, white arrows**). In the post-rugal field, NFH+ and NFH-afferents were identified coursing up into lamina propria invaginations towards the epithelium (**Figure 6c, blue arrow**). Merkel cells were also found in clusters at the base of epithelial pegs (**Figure 6c, white arrows**). A higher magnification view of a Merkel cell cluster shows Merkel cells contacted with NFH+ afferents (**Figure 6d, white arrows**), and some Merkel cells without obvious neuronal contacts (**Figure 6d, blue arrows**). At the tips of lamina propria invaginations of hard palate rugae, NFH+ afferents coalesced into endings that appeared to be corpuscular (**Figure 7a, white arrows**). Larger NFH+ terminal fields, reminiscent of glomerular endings identified in mice and donated specimens were also identified in hard palate rugae of healthy subjects (**Figure 7b, white arrow**). In the post-rugal field, Meissner’s corpuscles were identified in lamina propria invaginations (**Figure 7c, white arrow**). Ultraterminals were also occasionally found in the post-rugal field (**Figure 7d, white arrow**). Overall, these data demonstrate a similar arrangement of somatosensory endings between the rugae and post-rugal field of the hard palate.

**Figure 6.**
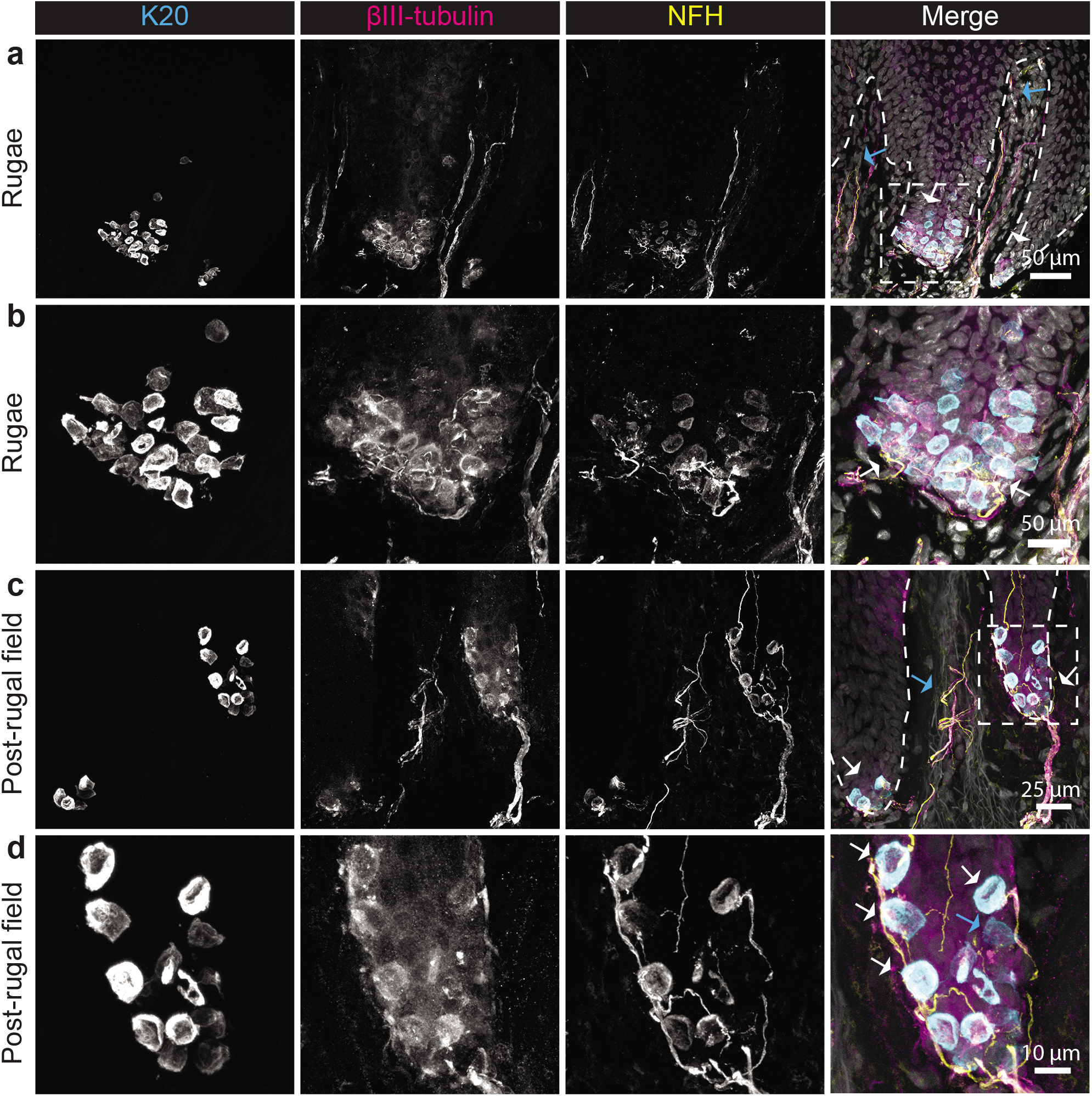
Clusters of Merkel cell-neurite complexes line the base of epithelial pegs of the hard palate rugae and post rugal field. Hard palate biopsies from healthy donors were sectioned and stained with markers for Merkel cells (K20, left column), all neurons (βIII-tubulin, second column), and myelinated neurons (NFH, third column). Epithelial-lamina propria boundary is demarcated by dashed lines. **a.** In rugae, the base of epithelial pegs were densely populated by Merkel cell clusters, visualized by K20 immunoreactivity (white arrows). Myelinated neurons visualized by NFH+ afferents extended into apical portions of lamina propria (blue arrows **b.** Higher magnification view of Merkel cell cluster shown in **a** (dotted box). NFH+ neuronal afferents innervated epithelia (white arrow). **c.** The base of epithelial pegs of the post-rugal field was also densely packed with Merkel cells (white arrows). NFH+βIII+ afferents also extended into the apical regions of the lamina propria (blue arrows). **d.** Higher magnification view of Merkel cell cluster in **c.** (dotted box). Merkel cells in this cluster were innervated by NFH+ afferents (white arrows). Some Merkel cells were not contacted by afferents (blue arrow).

**Figure 7.**
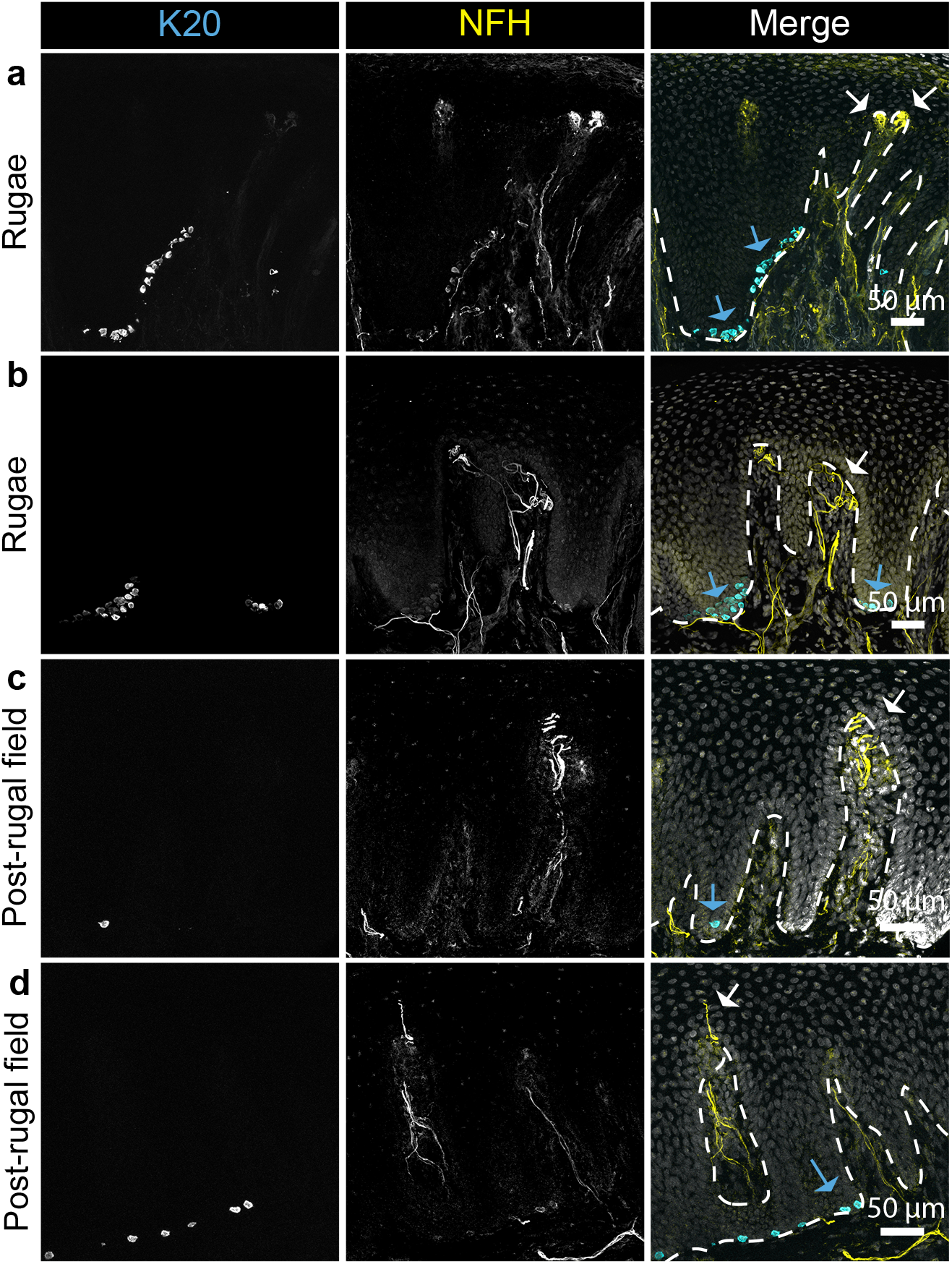
Corpuscles and ultraterminals are located in the hard palate epithelium. Hard palate biopsies from healthy donors were sectioned and stained with markers for Merkel cells (K20, left column and myelinated neurons (NFH, middle column). Epithelial-lamina propria boundary is demarcated by dashed lines. **a.** Meissner corpuscles were located in rugae of hard palate epithelium (white arrows). Merkel cell clusters were found along base of epithelium (blue arrows). **b.** Glomerular endings were identified in apical regions of rugae epithelial pegs (white arrow). **c.** In the post-rugal field, Meissner corpuscles were located in the apical portions of epithelial pegs (white arrows); sparse Merkel cells were located at the base of epithelium (blue arrows). **d.** Occasional ultraterminals were identified in post-rugal field (white arrow).

## Discussion

In previous studies, oral mechanoreceptors have often been lumped into one group, rather than considering the contributions of specialized classes. To achieve a better understanding of the diversity and distribution of oral mechanosensory neurons, we surveyed oral sensory neurons in human tongue and hard palate to make predictions about functionality. This report describes the distribution of sensory innervation of human oral epithelia derived from healthy donors and donated specimens (**Figure 8)**. We found that hard palate was innervated with Meissner’s corpuscles and glomerular endings and that hard palate epithelia were lined with Merkel cell-neurite complexes. In the tongue, fungiform papillae showed occasional Meissner’s corpuscles. Myelinated afferents also surrounded the taste buds and innervated the epithelium adjacent to taste buds. Filiform papillae had abundant end bulbs of Krause as well as free endings in the lamina propria. Occasional K20+ putative Merkel cells were found in the epithelia of filiform papillae as well as densities of myelinated endings that aggregated at the epithelial border. Thus, we conclude that the general organization of tongue and hard palate innervation are conserved between humans and mice, as described in previous studies (Moayedi et al., 2018), with some additional afferent types found in the human tongue.

**Figure 8.**
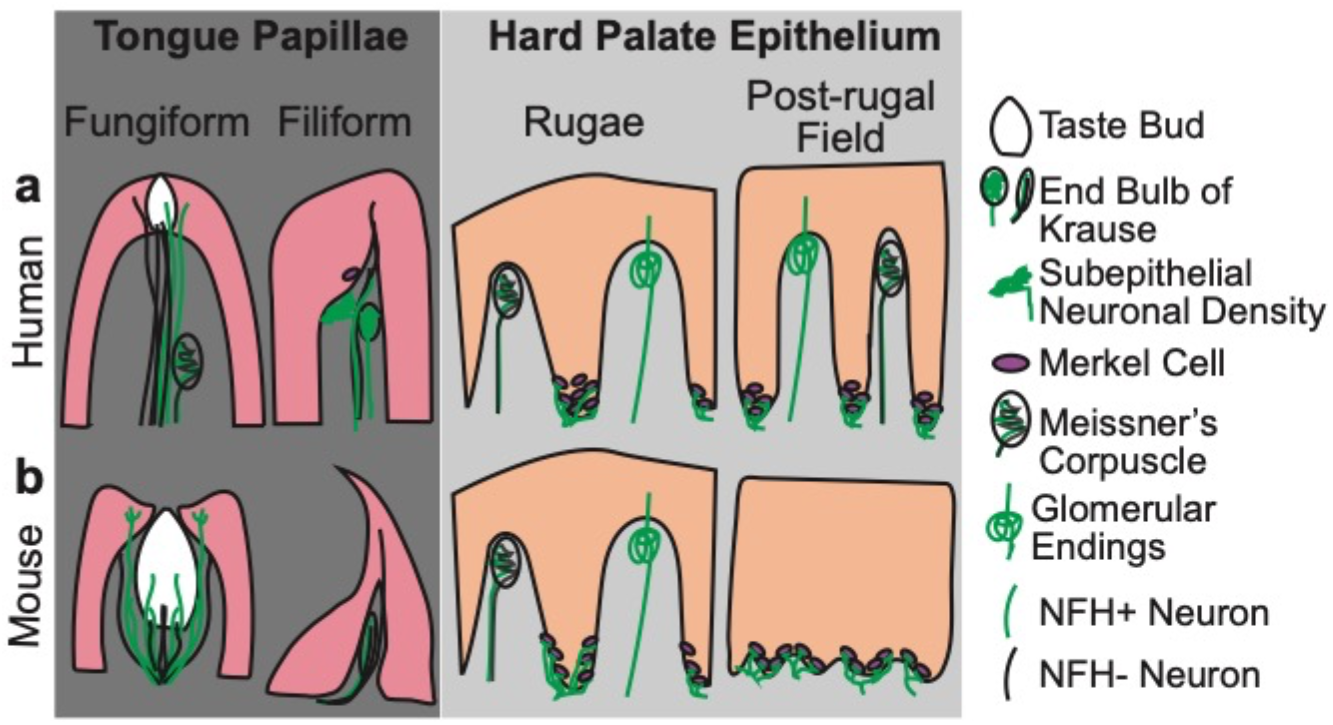
Comparison between somatosensory innervation of human and murine oral epithelia. **a.** Schematic showing somatosensory innervation of the tongue and hard palate epithelium of humans. **b.** Comparable structures in mouse tongue and hard palate.

Mechanosensation in the oral cavity is thought to subserve a range of essential functions including speech production, flavor sensation, and feeding biomechanics. Mechanosensory interactions between the tongue and hard palate are important for positioning the tongue during speech articulation (Niemi et al., 2002; Palmer, 1979). Loss of contact between the tongue and hard palate through application of dentures or loss of sensation can cause deficits in speech articulation. In regards to flavor, mechanosensation through the tongue, hard palate, teeth, and cheeks is thought to convey a variety of textural features of foods including grittiness, stickiness, viscosity, hardness, moisture content, and greasiness. Mechanoreceptors in the hard palate are particularly important for sensing thickness of food (Kutter, 2011). In addition to these features, mechanosensation is also thought to contribute to sensations of astringency, fat, mouthfeel, and flavor binding (Green, 2003; Haggard & de Boer, 2014; Linne & Simons, 2017; Liu et al., 2016). Finally, force sensation in the teeth and bolus characteristics sensed through the tongue and hard palate are believed to be important factors influencing feeding biomechanics (Inoue et al., 1989; Moore et al., 2014; Thexton et al., 1980; Travers & Norgren, 1986). Despite the importance of mechanosensation in oral functions, we have a limited knowledge of the identities and functions of types of mechanosensory neurons present in oral tissues.

As mechanosensation is thought to underlie oral functions such as feeding and speech, differences in the density and distribution of mechanosensory afferents between individuals may affect these functions. For example, PROP tasters and supertasters have enhanced tongue stereognosis as well as differences in flavor discrimination compared to non-tasters (Bangcuyo & Simons, 2017; Essick et al., 2003; Tepper & Nurse, 1997). Structurally, PROP tasters have a higher density of fungiform taste papillae compared to non-tasters and are postulated to have a correlated increase in density of trigeminal innervation to fungiform papillae (Bartoshuk et al., 1994). Yet, it has yet to be tested whether an increased density or altered distribution of mechanosensory afferents are correlated with increased fungiform papilla density. Similarly, mechanosensory innervation and function could affect speech abilities, thus further investigations into alterations in mechanosensory function in individuals with speech pathologies are warranted (McNutt, 1977; Niemi et al., 2006; Niemi et al., 2002; Petrosino et al., 1987). Furthermore, oral stereognosis and discrimination abilities decline with aging, which have potential impacts on feeding and speech abilities (Bangcuyo & Simons, 2017; Park, 2017). Additional studies are needed to see if there are correlated decreases in density or distribution of mechanosensory end organs in humans, as occurs in mice (Ikebe et al., 2007; Moayedi et al., 2018). Understanding the distribution of oral mechanoreceptors in healthy tissue provides a basis for further functional comparisons.

The tissue in which mechanosensory end organs are embedded potentially impacts the sensitivity of those endings to stress and strain, thus is important to consider when postulating function. For example, modeling approaches suggest that the topology of the tongue enhances the strains experienced in the tongue (Lauga et al., 2016). The bases of filiform papillae are calculated to experience maximal strain in modeling studies, suggesting there is an evolutionary advantage to positioning mechanoreceptors in this region for detecting strains, such as those experienced when detecting flow or shear. The presence of putative mechanosensory endings in this region of the filiform, including end bulbs of Krause, suggests that mechanoreceptors in filiform are optimally positioned to detect deflections in filiform papillae. By contrast, the hard palate rugae have much more rigid structures where mechanosensory innervation is found both at the apex and base of epithelial pegs. The ridged structures may help to increase frictional forces experienced by mechanosensory neurons in both the tongue and hard palate during speech and feeding. The importance of structural interactions is evidenced by the fact that denture wearers experience significant improvement in speech when the maxillary denture is fashioned with artificial rugae (Adaki et al., 2013; Zaki Mahross & Baroudi, 2015). Further exploration is warranted to address the roles of palatal epithelia and mechanoreceptors embedded within them in aiding in speech and feeding. Keratinization can also alter tissue mechanics and dampen sensation, as is the case in callouses, we would expect this factor to alter the forces transduced by mechanosensory endings. Future functional studies should take these variables into account.

### Tongue innervation

Few previous reports have investigated the anatomical diversity of human tongue sensory innervation. An early study described free nerve endings and mucocutaneous end organs in tongue papillae; however, the authors concluded that these tongue sensory endings lacked anatomical specializations (Marlow et al., 1965). Subsequent studies have demonstrated that the human tongue is innervated by physiologically distinct mechanosensory subtypes, and shown examples of innervation surrounding taste buds in fungiform papilla (Hilliges et al., 1996). By contrast, studies of tongue innervation in rodents, cats and nonhuman primates indicate that that the mammalian tongue is generally innervated by a number of structurally distinct end organs (Moayedi et al., 2018; Munger, 1973; Spassova, 1974; Suemune et al., 1992; Toyoshima et al., 1987; Zahm & Munger, 1985).

To resolve these discrepancies, we analyzed structure and innervation of human fungiform papillae and compared results to previous findings in mice (**Figure 8**; (Moayedi et al., 2018). Mouse and human fungiform papillae showed several structural differences: mouse papillae are small with a single taste bud in each papilla, whereas humans have large papillae with multiple taste buds on the surface. In mouse, dermal invaginations surrounds the fungiform taste bud, with neuronal innervation wrapping around the bud in these invaginations and extending into the overlying epidermis. Importantly, these neurons surrounding the taste bud were immunoreactive for the mechanosensory ion channel Piezo2, indicating that they are mechanoreceptors (Moayedi et al., 2018). Human fungiform papillae taste buds did not appear to have these invaginations and the corresponding innervation. Instead, human taste buds had a dense network of innervation in the lamina propria below the taste bud. Moreover, observed in mice, NFH+ and NFH-fibers extended into the taste bud and into the epithelia surrounding the taste bud. NFH-afferents in this region are likely a composed of both taste afferents and unmyelinated c-fibers involved in transduction of thermal, chemical, and noxious stimuli. We hypothesize that the NFH+ fibers surrounding the taste bud include mechanoreceptors that are analogous to the Piezo2+ endings identified surrounding taste cells in mice (Donnelly et al., 2018; Moayedi et al., 2018). In addition to putative mechanosensory endings surrounding taste buds, we found Meissner’s corpuscles in a human fungiform papilla. To our knowledge, Meissner’s corpuscles have not been previously found in human fungiform papillae, although they have been described in non-human primates (Zahm & Munger, 1985). These corpuscular endings are important for detecting vibrations and could play an important role in providing sensory feedback during speech, particularly in generating sounds where the tip of the tongue, where fungiform papillae are most dense, is in contact with the hard palate or teeth (Niemi et al., 2006; Niemi et al., 2002).

This arrangement of mechanoreceptors in fungiform papillae suggests that having taste and touch in close contact serves an important role in integrating sensations. This colocalization could aid in taste localization on the tongue (Todrank & Bartoshuk, 1991). This ability is thought to be an important factor in binding the sensations of olfaction, taste, and touch into a unified flavor, and generalizing taste location across the oral cavity (discussed in (Green, 2003). Future studies are needed to investigate whether particular mechanoreceptors or types of mechanical stimuli (e.g. moving vs. pressure) are required for this phenomenon.

We next analyzed morphology and innervation of human filiform papillae. Similar to mouse, human filiform papillae were innervated by encapsulated putative mechanoreceptors and had innervation extending into the apical papillae (**Figure 8**; (Moayedi et al., 2018). Human filiform papillae also had several notable differences in structure from those of mouse. In humans, filiform papillae had flatter peaks compared to mouse, with wider lamina propria core regions. These differences in macroscopic structures are likely due to different functions of the tongue. In addition to flavor assessment, the mouse tongue is specialized for grooming, which is aided by the rough keratinized ‘combs’ created by the array of filiform papillae. The human tongue, on the other hand, is more adept for speech and does not necessitate grooming adaptations. In mice, the filiform cores are innervated with a single bundle of fibers surrounded by Nestin+ cells, collectively termed end bulbs of Krause. These mouse afferent bundles contain both NFH+ and NFH-neuronal fibers and expressed the mechanosensory ion channel PIEZO2 (Coste et al., 2010; Moayedi et al., 2018; Ranade et al., 2014). In humans, a single filiform papilla contained several end bulbs of Krause with NFH+ and NFH-fibers. Human end bulbs had more spherical structures than those found in mice. We hypothesize that these end-organs are a major contributor to mechanosensation in filiform papillae. Small bundles of fibers traversing into the apical lamina propria invaginations containing NFH-fibers were also found, as described previously in mice. These fibers are likely c-fibers that detect temperature changes or chemical agonists. Two types of NFH+ endings that were not previously defined in the literature were also identified. First, bundles of NFH+ afferents terminating just below the epithelium in humans were termed subepithelial neuronal densities. These endings are similar to disk-like mucocutaneous terminals described by Marlow and colleagues (Marlow et al., 1965). Second, NFH+ endings were found to innervate the basal epithelium of filiform papillae. The relevance of these afferents to somatosensation is not known. Overall, this innervation pattern is more complex than that of the mouse, suggesting that humans experience a richer repertoire of mechanical sensations when filiform papillae are stimulated.

Unlike mouse, human filiform papilla also contained occasional K20+ cells that resemble Merkel cells. Unexpectedly, these K20+ cells were not associated with innervation. These K20+ cells might be another cell type that expresses this keratin, or they might be Merkel cells that perform a paracrine role in the epithelium (Xiao et al., 2014). As previously observed in the mouse tongue, these results indicate that Merkel cells are not likely contributors to pressure sensation in the human tongue, as they are sparse in density and have no apparent contacts with peripheral neurons (Moayedi et al., 2018).

Four physiologically distinct types of oral mechanoreceptors have been identified in human tongues (Trulsson & Essick, 1997, 2010). These include one rapidly adapting (RA) response signature, two types of slowly adapting (SAI and SAII) neurons, and one class of deep mechanoreceptive afferent presumed to be proprioceptors. In studies in the glabrous skin, RA responses typically indicate activation of an encapsulated end-organ such as Meissner’s or Pacinian corpuscles, SAI responses signify Merkel cell-neurite complex activation, while SAII responses are attributed to Ruffini endings (Bautista & Lumpkin, 2011; Johnson et al., 2000). The neuronal afferents from which these physiological signatures derive from in the tongue has not been tested directly. The RA response type is likely to be derived from encapsulated Meissner’s corpuscles and potentially the end bulbs of Krause in the human tongue (Sakada, 1983). The origins of the two SA response categories in the tongue are unclear. The SAI response is typically derived from Merkel-cell neurite complexes. Although we did identify sparse K20+ cells in the filiform papillae, we note that these were uncommon and did not appear to have contacts with neurons, thus are not likely the origin of the SAI response. This suggests that the SAI response may be derived from a different end organ in the tongue. Ruffini endings are hypothesized to be the origin of the SAII response, although we did not identify these ending types in human oral tissue this study does not eliminate that possibility. Future studies should identify the physiological responses of the anatomical classes identified this study, including subepithelial neuronal densities and NFH+ neurons surrounding the fungiform papillae, to test the hypothesis that these mediate slowly adapting responses in the human tongue.

The density and composition of mechanoreceptors in fungiform and filiform papillae provides the biological basis for mechanosensory qualities that they are tuned to detect, and thus provides hints as to the functions of mechanoreceptors within them. Previous observations suggest that increased fungiform papillae density is correlated with increased tactile acuity in a stereognostic tongue letter detection task (Bangcuyo & Simons, 2017; Essick et al., 2003). This suggests that mechanoreceptors in fungiform papillae are particularly tuned to detect edges, an object feature that is attributed to Merkel cells-neurite complexes via the SAI response (Johnson & Lamb, 1981). We found no evidence of Merkel cells within the fungiform papillae, and no evidence of innervated Merkel cells within the tongue. Rather, we found evidence of at least two putative mechanosensory afferent types in the human fungiform papillae: Meissner’s corpuscles and myelinated afferents surrounding the taste buds. The former is known to give an RA-type response, raising the possibility that the latter could generate an SA response. This hypothesis should be addressed in future studies. Interestingly, ability to detect surface roughness seems to be independent of fungiform density, raising this possibility that filiform mechanoreceptors may play an important role in this feature of food texture assessment (Linne & Simons, 2017). As RA type responses are important for encoding texture, this suggests that the end bulbs of Krause that are the most abundant end organ in filiform papillae may be an RA type mechanoreceptor (Lamb, 1983). This possibility should be addressed in future studies.

### Hard palate innervation

Human hard palate rugae were found to have similar morphology and innervation to that of mice, suggesting that innervation patterns of rodent and human hard palates are highly homologous (**Figure 8**; (Moayedi et al., 2018; Nunzi et al., 2004). The hard palate was lined with epithelial pegs innervated by NFH+ and NFH-afferents. These afferents coalesced to form corpuscular structures or glomerular endings. Merkel cells densely lined the bases of the epithelial pegs, as previously found in mice and humans (Gairns, 1954; Moayedi et al., 2018; Nunzi et al., 2004). Merkel cells in human palate were often innervated by NFH+ afferents, forming Merkel cell-neurite complexes. Collectively, the hard palate rugae in both mouse and humans are homologous in neuronal architecture and composition to other regions of high tactile acuity, such as those of the rete ridges of the fingertips (Cauna, 1954). These share a common arrangement of Merkel cells at the base of epithelial pegs and Meissner’s corpuscles jutting between pegs to the apical regions of lamina propria. We hypothesize that this arrangement confers the palate with high tactile acuity that is essential for speech and flavor construction.

The morphology of the post-rugal field was strikingly different from that identified in mice. Human tissue contained many epithelial pegs that were innervated by NFH+ fibers with occasional ultra-terminals and Meissner’s corpuscles with dense clusters of Merkel cells at the base. Mice did not have epithelial pegs or ultra-terminals in this region, rather they had a thickened epidermal layer with cluster of Merkel cells at the epidermal-lamina propria junction (Moayedi et al., 2018). Thus, in humans, innervation of the back of the hard palate more closely mirrored that of palate rugae. As humans have a different distribution of hard palate rugae compared to back of the hard palate, future studies should be aimed at defining whether the different regions of this area have uniform innervation. This arrangement of primarily Meissner’s corpuscles and dense Merkel cells in the human hard palate suggests that this region of the hard palate is tuned to detect sustained force, vibration, and textures. These percepts are particularly important for detection of mouthcoating and other flavor features, suggesting the importance of the palate in these processes.

### Summary

In summary, this study provides a systematic evaluation of neural structures of healthy human hard palate and tongue. We have confirmed previous descriptions of innervation of the human hard palate rugae, showing that it is innervated by Meissner’s corpuscles, glomerular endings, and Merkel cell-neurite complexes. We have extended these findings to the post-rugal field, which is primarily innervated by Meissner’s corpuscles and Merkel cell-neurite complexes. These palatal structures are well poised to contribute to high acuity tactile perception needed for feeding and speech. In the tongue, this analysis has identified several novel afferent endings in fungiform and filiform papillae. The fungiform papillae are innervated by occasional Meissner’s corpuscles, and their epithelial layers are innervated by terminals of myelinated afferents. Filiform papillae contain a high density of end bulbs of Krause, subepithelial neuronal densities, and NFH+ neurons innervating the epithelium. The functional contributions of these tongue mechanoreceptors to oral sensations are currently unknown and require further exploration. Collectively, these somatosensory endings provide a structural basis for the rich mechanosensory abilities of human oral tissues.

## Authorship

All authors had full access to the data in the study, and take responsibility for data integrity and the accuracy of data analysis.

Conceptualization: YM, SM, EAL; Methodology: YM, EAL, MP, AK; Formal Analysis: YM, EAL; Investigation: YM, MP; Resources: EAL; Data Curation: YM, EAL; Writing-Original Draft: YM; Writing-Review and Editing: YM, MP, AK, SM, EAL; Visualization: YM; Supervision: EAL; Project Administration: EAL; Funding Acquisition: EAL

## Acknowledgments

We thank Drs. Paulette Bernd and Michael Pitman for facilitating collection of anatomical donor specimens and Dr. Benjamin Le Reverend for impactful discussions during conceptualization of this project. This work was supported by SPN, SA; Berrie Foundation Initiative on the Neurobiology of Obesity (EAL, YM); Thompson Family Foundation Initiative on Chemotherapy Induced Peripheral Neuropathy and Sensory Neuroscience (YM); and NIH NIAMS R01AR051219 (EAL). Imaging was performed with support from the Herbert Irving Comprehensive Cancer Center Confocal and Specialized Microscopy Shared Resources (P30CA013696) and Zuckerman Institute’s Cellular Imaging Platform.

## Conflict of interest statement

SM is an employee of Sociétés des Produits Nestlé S.A.

## Notes

### Competing Interest Statement

SM is an employee of Societes des Produits Nestle S.A.

